# The threshold of binocularity: natural image statistics explain the reduction of visual acuity in peripheral vision

**DOI:** 10.1101/131177

**Authors:** David W. Hunter, Paul B. Hibbard

## Abstract

Visual acuity is greatest in the centre of the visual field, peaking in the fovea and degrading significantly towards the periphery. The rate of decay of visual performance with eccentricity depends strongly on the stimuli and task used in measurement. While detailed measures of this decay have been made across a broad range of tasks, a comprehensive theoretical account of this phenomenon is lacking. We demonstrate that the decay in visual performance can be attributed to the efficient encoding of binocular information in natural scenes. The efficient coding hypothesis holds that the early stages of visual processing attempt to form an efficient coding of ecologically valid stimuli. Using Independent Component Analysis to learn an efficient coding of stereoscopic images, we show that the ratio of binocular to monocular components varied with eccentricity at the same rate as human stereo acuity and Vernier acuity. Our results demonstrate that the organisation of the visual cortex is dependent on the underlying statistics of binocular scenes and, strikingly, that monocular acuity depends on the mechanisms by which the visual cortex processes binocular information. This result has important theoretical implications for understanding the encoding of visual information in the brain.

## Introduction

It has long been known that visual acuity is greatest in the centre of the visual field, peaking in the fovea and degrading significantly towards the periphery [1], [14]. For very simple tasks, such as detection of low contrast targets, the rate of change of the cortical magnification factor is correlated with the rate of change of task performance [26]. Indeed, substantial evidence has been provided to link performance with the physiology of the human visual system; our fundamental visual acuity [31] and sensitivity to differences in orientation [4], and spatial frequency [25, 26], are all linked to the density of retinal ganglion cells.

Detailed psychophysical measurements of these performance differences are of great theoretical importance as they identify informational bottlenecks in visual processing [15]. If Vernier acuity for example is limited, then any task dependent on the same information will be limited to the same degree [15]. This theoretical insight is particularly revealing given that the effects of eccentricity can be directly related to physiological properties such as the density of retinal ganglion cells (for simple tasks) [25, 26] or the organisation of ocular dominance columns (in the case of Vernier acuity and crowding) [15].

In binocular vision, human depth acuity is substantially poorer in eccentric regions than predicted by cortical magnification [9], however the rate of decay is similar to that for Vernier Acuity [27, 29]. The decay rate is also dependent on the spatial frequency of the stimulus, with feature detection substantially less accurate at eccentric locations for fine scale features than for coarse-scale features [27].

These physiological and psychophysical measures provide insight into the mechanisms of early vision, however understanding why the visual system works the way it does requires a different approach. The visual scene contains many redundancies that can be exploited in order to process stimuli. Barlow proposed that early stages of the visual system (retina-LGN-V1) form an efficient coding of ecologically valid stimuli [3]. This idea was further developed by Marr who proposed a visual hierarchy of neurons specialised to detect particular specific visual patterns depending on the relative frequency of their occurrence [19].

Olshausen and Field showed that Independent Component Analysis applied to inputs from photographs produced filters with a similar structure to the receptive fields of simple cells in V1 [22]. Similar results were found for binocular images by Hoyer and Hyvärinen [11]. As with simple-cell receptive fields, ICA components can be characterised in terms of their position, frequency, phase, orientation and binocular properties. The distributions of ICA component characteristics share substantial similarities with the distributions of V1 receptive fields, in particular the distributions of horizontal and vertical disparities closely resemble the distributions of horizontal and vertical disparities in V1 [12].

We propose that the requirement for binocular vision imposes a constraint on the finest spatial scale at which visual processing can reliably occur. This threshold of binocularity constraint can then account for both the reduction of binocular acuity with eccentricity and stimulus wavelength, and also the reduction in acuity for purely monocular tasks. As such, the need for binocular vision can be seen as imposing a fundamental bottleneck on visual encoding which extends beyond the requirements of binocular depth perception, and provides a direct explanation for the overall fall-off in visual acuity with eccentricity.

### Threshold of Binocularity

Central to our theory is the idea that some visual features are optimally encoded binocularly, resulting in binocularly tuned cells. Conversely, other features are optimally encoded monocularly leading to cells the respond maximally to features in one eye only. Typically, ICA will learn both monocular and binocular components [11, 12]. In both cases, this reflects the redundancy present in natural binocular images. As eccentricity increases, the distribution of binocular disparities increases [10, 18, 23, 28] thus reducing the similarity expected in a corresponding region of an image pair across the two eyes.

For a given image eccentricity, the degree to which binocular processing is possible will depend on the availability of a sufficient number of binocular components. Since this will depend on the spatial scale of analysis, we can define a Threshold of Binocularity: the smallest spatial scale at which sufficient binocular components exist to support binocular processing. Given the dependence of binocular components on both eccentricity and scale, the Threshold of Binocularity will increase with eccentricity. It follows from this that binocular processing will then only be possible at increasingly coarse spatial scales as eccentricity increases. Fine scale features in eccentric regions that cannot be binocularly matched would need to be processed monocularly and would not (at least directly) form part of a cyclopean percept. The Threshold Angle of Binocularity thus poses a constraint on the highest spatial scale of processing, at each image eccentricity, that is consistent with binocular integration. Critically, if we wish to ensure binocular vision across the visual field, then it follows that both monocular and binocular processing will be constrained to only occur at progressively coarser scales with increasing eccentricity.

## Results

We generated an efficient coding of stereoscopic images using Independent Component Analysis. The proportion of binocular components varied substantially depending on both the angle of eccentricity and the wavelength of the feature, as shown in Figure 1. The proportion of binocular components is close to zero at short wavelengths and large eccentricities, and close to 1 at long wavelengths and small eccentricities. This arises because long wavelength features are more likely to overlap in each view and thus merge into a single binocular feature. Similarly, since the range of disparities is greater at larger eccentricities than in the fovea [10, 28], we would expect a lower proportion of binocular components in these areas. Between these two points a clear and rapid transition occurs from predominantly monocular components to predominantly binocular components. We defined the threshold of binocularity as the iso-contour where 50% of components were binocular and 50% monocular. This threshold was estimated be linear interpolation and is shown as a black line in figure 1. We used the iso-contour of this transition to calculate the rate of decay with eccentricity, *E*_2_ which we found to be 0.7421. This is high similar to the human stereo-acuity results presented by Fendick and Westheimer of *E*_2_= 0.81 ([9] as reported by [29]).

**Figure 1.**
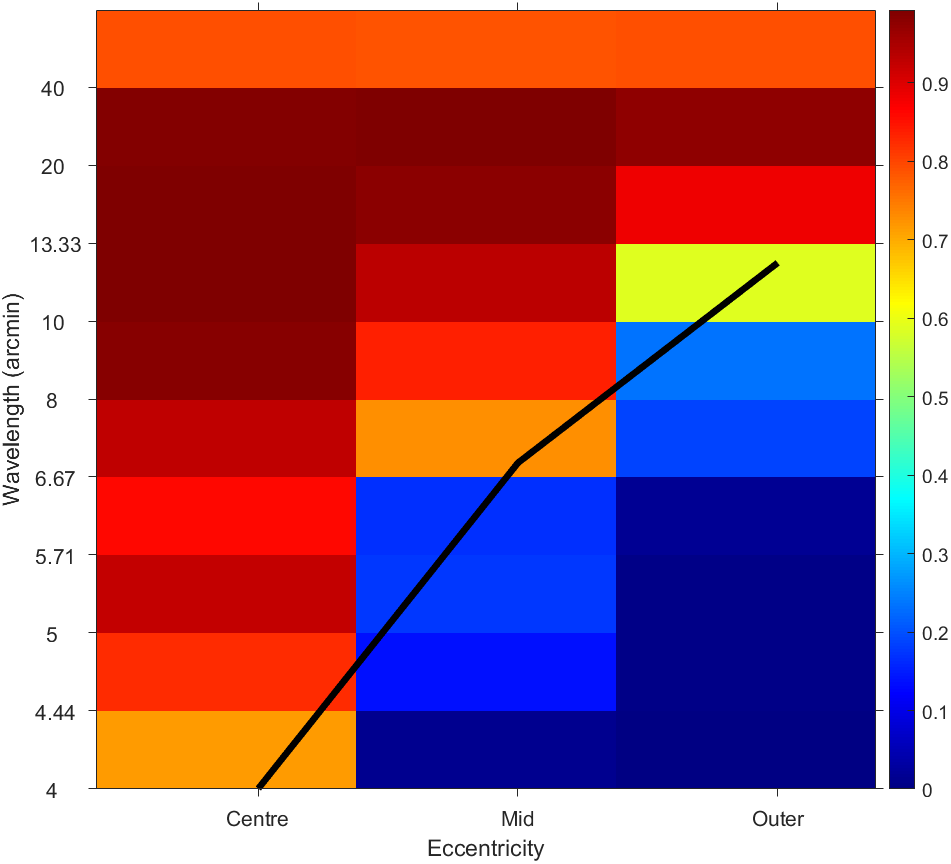
2D histogram of the proportion of binocular components across wavelength and eccentricity. Figure 1. Red shows 100% binocular, while blue shows 0% binocular components in each wavelength/eccentricity bin. The black line shows the Threshold of Binocularity (50% binocular components by linear interpolation).

### A wavelength-eccentricity surface on binocular thresholds

*E*_2_ captures a linear effect of eccentricity in one dimension only. In the previous analysis, *E*_2_ was used to describe the effect of eccentricity on the wavelength threshold: the spatial scale up to which binocular processing can occur, given the components learned through ICA. An alternative approach is to analyse the impact of both eccentricity and scale on the ratio of binocular to monocular components. In this approach *E*_2_ is calculated for each wavelength by measuring the slope of the binocular/monocular ratio, i.e. the rate of performance decline for each wavelength. A 2D sigmoid function was fitted to the data as a smoothing function and *E*_2_ was calculated as the eccentricity at which the ratio of binocular/monocular components was half of that of the fovea. The fitted sigmoid function is plotted in Figure 1. The iso-contour showing the binocular threshold hold (at 50% binocular) is shown in black, this is the omnibus *E*_2_ from the previous experiment. *E*_2_ for each wavelength is calculated on vertical slices of the 2D sigmoid take at each wavelength.

Values of *E*_2_ for each wavelength are shown in Table 1. Results reported for human stereo-acuity by Siderov and Harwerth 9 are shown where there is appropriate data. It is worth noting that our estimate of *E*_2_ matches that of Siderov and Harwerth at 7.5 c/arcmin however only one wavelength from their data lies within the reliable range of our data. The second comparison at 30 arcmin is within the standard error of the result quoted by Siderov and Harwerth however our value is outside the range of our results and has been imputed.

**Table 1.** *E*_2_ and Binocular/monocular threshold values from fitted sigmoid in figure 2 for selected wavelengths. See main text for details of calculation. *-Values outside of the range of the data are not reliable.

| Wavelength (arcmin) | Threshold (arcmin) | $E_2$ | $E_2$ Siderov and Harwerth 1994 (standard deviation in brackets) |
| --- | --- | --- | --- |
| 4 | 1.77622 | 1.73133 |  |
| 5 | 2.007294 | 2.449983 |  |
| 6 | 2.836966 | 3.238440 |  |
| 7.5 | 4.081473 | 4.081473 | 3.8 (0.6) |
| 9 | 5.325981 | 5.706599 |  |
| 13 | 8.644669 | 9.024467 |  |
| 27 | 20.260074 | 20.639863 |  |
| 30 | 22.749090 | 23.128878 | 13.4 (9.5) |

**Figure 2.**
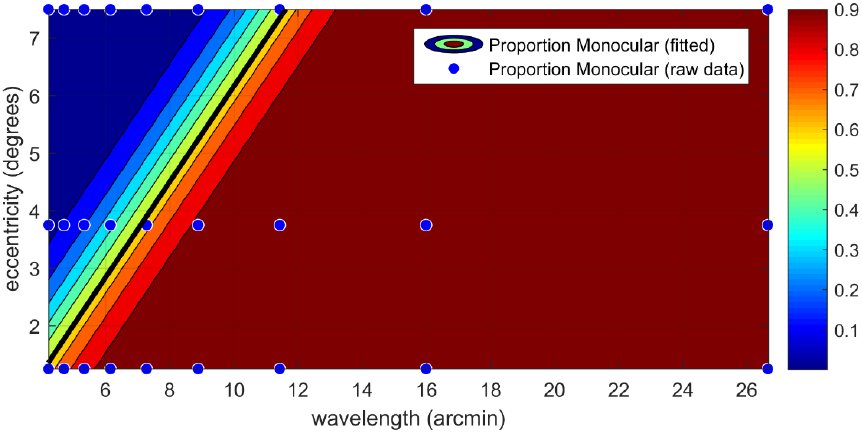
Proportion of binocular components across wavelength and eccentricity. The heatmap shows the iso-contours of a sigmoid fitted to the datapoints (blue marks). The iso-contour showing 50% binocular is highlighted in black.

### Comparing our predictions with human obervers

Our ICA results allow us to predict *E*_2_, the rate of decay in stereoscopic visual acuity with eccentricity. In order to validate our prediction, we compared our results with measurements of *E*_2_ in humans obtained psychophysically. Figure 3 shows a log-log plot of *E*_2_ as estimated from the fitted 2D psychometric function together with data from Siderov and Harwerth [27] table 3. Siderov and Harwerth gathered data from two observers; both are plotted in figure 3 together with the mean across the two. *E*_2_ was estimated from the 2D psychometric function (Table 1) and plotted as a black line. As the frequencies of the two plots do not overlap the psychometric measurements of Siderov and Harwerth were interpolated over the range of ICA frequencies using linear regression. ICA data was interpolated over the range of Siderov and Harwerth by sampling from the fitted psychometric function as wavelengths greater than those measured using ICA. Both ICA predictions and psychometric measurements match closely, with all ICA predictions lying within one standard deviation of the average psychophysical measurements. ICA predictions also closely match the linearly interpolated average from Siderov and Harwerth. The log-log gradient of Siderov and Harwerth was close to unity (-0.95), showing a linear relationship between frequency and *E*_2_, the gradient of our ICA data was -1.265, this is slightly surprising given the linear relationship between wavelength and eccentricity in our psychometric function, this effect is due to the proportion of binocular components being less than 100% at zero eccentricity. We do not measure the binocular/monocular ratio at zero eccentricity but rather from the area between zero and 150 arcmin.

**Figure 3.**
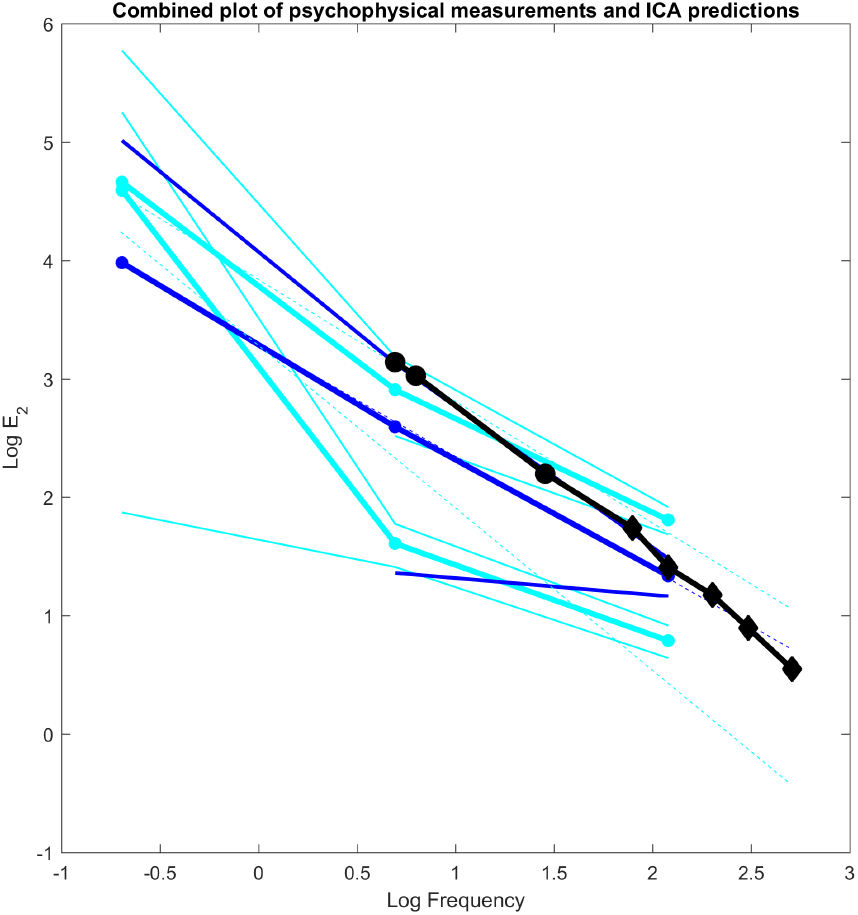
Combined log-log plot of psychophysical measures of *E*_2_ from 9 and ICA predications of *E*_2_. ICA predictions sampled from the fitted psychometric function are shown as a solid black line, diamonds mark samples taken from frequencies where the threshold is contained within the range of eccentricities in the original images (*≈*4 to 12 arcmin see figure 2), circles mark *E*_2_ values interpolated from the fitted 2D psychometric function. Data from [27] is shown as various shades of blue lines. Individual participant *E*_2_ are shown as cyan lines, the thick lines showing the measured *E*_2_, thin lines showing the first standard deviation, actual sample points are shown as a filled circle. The average across participants is shown as a dark blue line, the thick line showing the mean *E*_2_, thin dark blue lines showing the first standard deviation. The psychophysically measured functions were interpolated across the range of the ICA predictions by fitting a standard linear equation. Linearly interpolated lines are shown as dashed lines in the colour of the interpolated function.

## Discussion

Independent Component Analysis learns a sparse efficient linear coding of the binocular image space within a small portion of the image. Each component consists of a set of spatially close samples that vary linearly with respect to one another. A single binocular component forms a more efficient coding than two monocular components, so the appearance of monocular components indicate that in these regions of the binocular image space few correlations between left and right views exist.

There is a clear effect of wavelength on the proportion of binocular components that can be seen in figures 1 & 2. The proportions of binocular components and by extension the proportion of monocular components can be seen to approximate a sigmoid function from which we were able to calculate an approximation to the threshold between predominately binocular and predominately monocular components. A high prevalence of monocular components at a particular wavelength or eccentricity indicates that most low-level features vary independently from each other at that particular wavelength and location. It is not unreasonable to believe that humans would perform poorly in binocular tasks at eccentricities and wavelengths where monocular components predominate. Psychophysical measurement of binocular acuity by Siderov and Harwerth [27] show that this is indeed the case, with both lower binocular acuity and a steeper reduction in visual acuity with eccentricity for high frequency stimuli compared with low frequency stimuli Siderov and Harwerth [27].

We found that the binocular threshold wavelength varies in an approximately linear fashion with eccentricity, with an *E*_2_ of 0.74. In terms of frequency the binocular threshold function varies in the inverse exponential decay expected of an eccentricity function. This result places the rate of change of the binocular threshold squarely with the range of results of both binocular depth acuity [9, 27]. We have also shown that both the predicted values and general trend of our predications lie within one standard deviation of psychophysical measurements of *E*_2_ in humans [27]. This indicates that human binocular visual performance in eccentric regions is optimised to the statistics of binocular natural images.

The substantial reduction in monocular visual acuity with visual eccentricity has not yet been fully explained. Weymouth proposed retinal ganglion cells as the bottleneck on eccentric visual performance [31]. However, the greater loss of visual acuity in positional (e.g. Vernier) [16] and recognition [1] tasks verses simple motion detection [17] indicates that loss of fidelity occurs after retinal-ganglion cell computation.

Levi linked Vernier acuity to the structure of ocular dominance columns by measuring the range at which crowding affects Vernier acuity performance and comparing this retinotopically to the size of ocular dominance columns in V1. This indicates a physiological link between a purely monocular task and the binocular processing structure of the visual cortex. Binocularly tuned regions in V1 are physically located in close proximity to ocular dominance columns in layer 4 of V1 and are stimulated by feed-forward connections from these columns [21].

The value of *E*_2_ (0.74) measured here lies within the range of both binocular depth acuity (0.81 [9, 27] and monocular Vernier acuity (0.62-0.77) [16]. When put together with the physiological and psychophysical evidence that binocular depth acuity and monocular Vernier acuity are linked this suggests a strong link between the underlying statistical properties of binocular natural images and monocular visual acuity. This reflects a strong correlation between acuity thresholds for Vernier and stereoscopic tasks [20].

Formation of ocular dominance columns is widely viewed as being, at least in part, a response to the properties of visual stimuli [6]. Previously Chklovskii used wire-length minimisation to suggest a link between binocular disparity and ocular dominance columns [5]. The physiological evidence together with the psychophysical evidence indicate that both Vernier acuity and binocular acuity are limited by the same processes in V1.

We have found that the proportion of monocular to binocular components increases with eccentricity and this increase occurs at the same rate as both binocular depth acuity and monocular position acuity as measured in humans psychophysically. This indicates that binocular visual acuity is tuned to the statistics of natural images. Moreover, monocular visual acuity is linked to binocular acuity and therefore to the statistics of binocular images.

### Further thoughts

Binocular vision produces substantial advantages through enhanced signal to noise ratio, greater visual acuity and a direct perception of stereoscopic depth. Most of these benefits are found in the area of the fovea and toward the peripheries these benefits are greatly reduced. Adopting a binocular configuration is a trade-off between the added benefits in the fovea against reduced binocular fidelity in the peripheries.

## Materials and Methods

### The dataset

We used the binocular photographic image set of [10] as a source of natural images. The image-set consists of scenes covering a wide range of depths and disparities, from interior still life scenes on a light-table to outdoor scenes of woodland and beaches, taken around the town of St Andrews in Scotland, UK. The images were taken using a DSLR camera seated on a horizontal slide mount. Two images were taken of each scene, separated using the slide mount by 65mm, close to the average inter-pupillary separation for human adults [8]. In all scenes the cameras were independently verged and focused on an object in the centre of the image. This arrangement is intended to approximate human binocular vision.

### Preprocessing

Pairs of 25 by 25 pixel square patches were cut from the binocular images. Each pair of patches were cut from identical pixel locations within both left and right photographic images. Disparities in the scene are captured as small differences between the left and right rectangular image patches in each pair.

The raw patch images undergo a pre-processing stage in order to perform gain control and in order to meet the assumptions of the FastICA algorithm.

Given two raw vectorised images patches 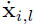 and 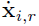 representing image patches cut from the left and right binocular images respectively, the vectors are centred by independently subtracting the mean from each vector. The vectors are independently normalised to remove the effects of local illumination.

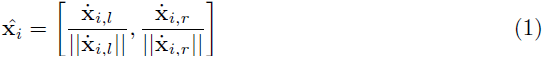

This function approximates gain control systems in early binocular features [7], reducing luminance differences between the two eyes and enhancing the phase differences that we are interested in. The FastICA algorithm requires normalised vectors,

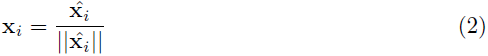

### Independent Component Analysis

We used Independent Component Analysis [13] on pairs of 25 by 25 pixel patches cut from identical locations relative to the top left of the image in both left and right views, this results in on square patch for each eye. As the sample location is fixed within the photographic frame binocular disparity within the scene is manifested as shift in location of the feature in one eye relative to another. The FastICA algorithm [13] attempts to learn a sparse linear decomposition of the dataset by maximising the Gaussianity of the loading (weights) matrix. If the underlying structure of the input dataset is linear-sparse, ICA will learn a sparse linear decomposition. The set of normalised image patches x_*i*_ form the matrix *X*. ICA decomposes *X* into two matrices, a factor matrix *W* and score matrix *A* as,

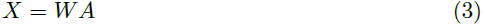

The columns of *W* are the independent components used in this analysis. The FastICA algorithm attempts to find *A* such that its elements are maximally non-Gaussian.

FastICA is a two stage process; a whitening pre-processing step using Principal Component Analysis and the ICA algorithm itself. Whitening using PCA is a necessary step to the FastICA algorithm [13] that acts as a low filter on the patch samples removing high frequency information. As high frequency information has a lower signal to noise ratio than low frequency information a low pass filter acts to reduce noise an increase the signal to noise ratio [2]. Together with the limits set by the patch size this stage produces a bandpass filtered representation of the binocular image data.

### 0.1 Eccentricity Regions

In order to examine the effects of eccentricity we restricted the sample area of the binocular image to a ring defined by angular eccentricity. Three regions were chosen; a 5*°*central region from a radius of 0 to 150 arcmin, a central region also covering 5*°*of visual arc from 150 arcmin to 300 arcmin radius and a 10*°*region from 300 to 600 arcmin. The total area of the visual field covered by the analysis was 20*°*in diameter. The sample regions are shown in Figure 4. Separate sets of patches were cut from the binocular image set from each of the three regions and separate sets of ICA component produced for each region.

**Figure 4.**
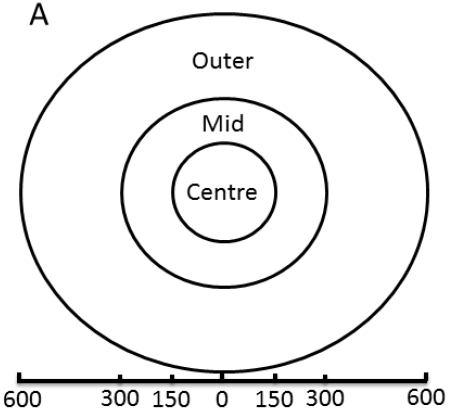
Eccentricity regions used in this analysis. Each region is defined by it’s angle of eccentricity in arcmins from the centre of the binocular image.

In total 3 groups of 100,000 25 by 25 by 2 pixel image patches were cut from each region in the binocular images. Each 25 by 25 pixel patch was cut from identical locations in both left and right images, as measured from the top-left edge of the image, and combined into a single binocular patch. Provided the disparities are smaller than the patch sizes, disparities between left and right images are retained within the cut patches. In order to capture a wider range of disparities and component frequencies than can be achieved within a 25 by 25 by 2 image patch we down-sampled the images at multiple scales and resampled the image regions. For each scale and image region we learned a set of sparse linear basis functions using Independent Components Analysis [13], resulting in 200 components (see figure 5 for examples).

**Figure 5.**
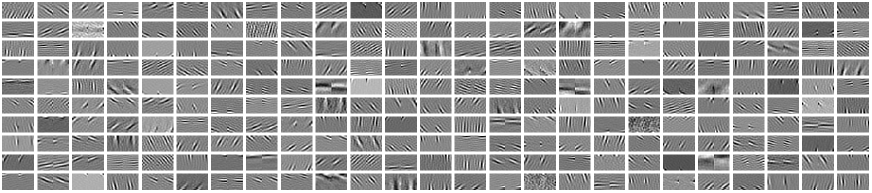
Examples of binocular linear components learned using Independent Component Analysis.

The entire process of image rescaling, patch sampling and ICA was repeated 10 times to produce 2000 components per region and scale combination. The linear components learned in this fashion resemble the receptive fields of simple-cells in V1 [11, 12, 22, 24]), allowing for comparison between the distribution ICA components learned from natural images and the distribution of known simple-cells in V1 [12, 24, 30]. The distribution of frequencies across all learned components is shown in figure 6. Components varied in the energy ratio between left and right eye parts, components with at least two thirds of energy in one eye were classified as monocular, components will a more equal energy distribution, at least one third of energy in each eye, were classified as binocular.

**Figure 6.**
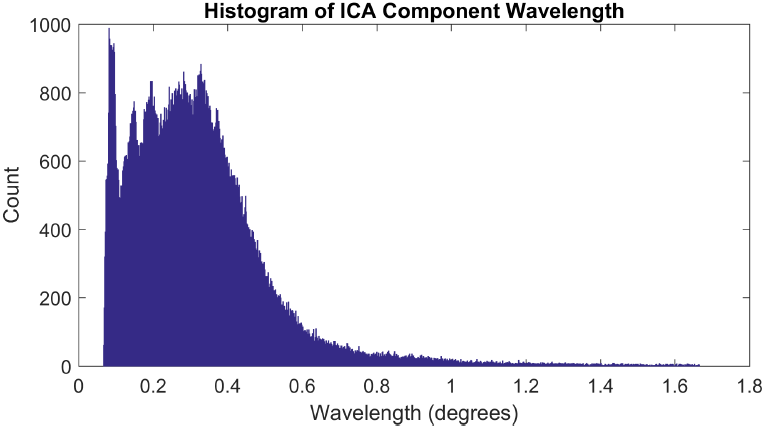
Histogram of the wavelengths of ICA components from all sample resolutions and eccentricities. The distribution of wavelengths for each individual sample resolution is not uniform resulting in a non-smooth distribution of wavelength. The local peaks show the separate maxima for each of the sample resolution sets.

### Cortical Magnification Factor

To explain this loss of acuity, the concept of the cortical magnification (M) was introduced by Rovamo et al. to measure the ratio between surface area of the visual cortex and the angle of visual field to which it is retinotopically mapped [26]. This factor varies across the visual field such that the inverse of M is roughly proportional to the angle of eccentricity. In a psychophysical context, the inverse *M*^-1^ represents a measure of sensitivity, such as the just noticeable difference between stimuli, or the minimum angle of resolution (MAR), depending on the experimental method and stimuli. It is assumed that visual sensitivity reflects the cortical resources that are devoted to representing visual information at each position in the image. *E*_2_ has the same interpretation in both contexts, allowing direct comparisons to be drawn between psychophysical and physiological measures. The Cortical Magnification Factor (*M*) is the ratio between the surface area of the visual cortex and the angle of visual field to which it is retinatopically mapped [26]. This factor varies across the visual field such that the inverse of M is roughly proportional to the angle of eccentricity. *M*^-1^ can therefore be described as:

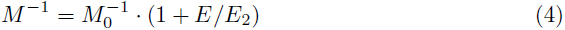

Where *E* is the eccentricity in degrees of visual arc, 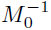 is the inverse cortical magnification factor (*°/mm*) at the fovea (*E* = 0*°*). *E*_2_ is the eccentricity at which the inverse cortical magnification factor is doubled relative to the fovea (or equivalently the cortical magnification factor is halved relative to the fovea). In a psychophysical context, *M*^-1^ represents a measure of sensitivity, such as the just noticeable difference between stimuli, or the minimum angle of resolution (MAR), depending on the experimental method and stimuli. It is assumed that visual sensitivity reflects the cortical resources that are devoted to representing visual information at each position in the image. *E*_2_ has the same interpretation in both contexts, allowing direct comparisons to be drawn between psychophysical and physiological measures.

### Threshold of Binocularity

We define the Threshold of Binocularity as the iso-contour where the ratio of binocular to monocular components generated by our learned model is 0.5 (see figure 1).

The rate of change of the Threshold of Binocularity can be measured in a similar manner to the Cortical Magnification Factor and the Minimum Angle of Resolution, using a slightly modified version of equation 4. Here *S*_0_ is the wavelength of the binocular/monocular threshold at an eccentricity of 0*°* and we replace MAR with the threshold of binocularity:

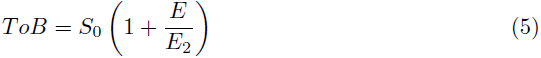

The values of *S*_0_ and *E*_2_ were determined from the data by using standard regression techniques, assuming that measurement errors were distributed normally. We found that *S*_0_ = -0.005*°* and *E*_2_ = -0.7421. As *S*_0_ indicates a negative wavelength at the fovea which is physically impossible, this results from interpolation from data-points in the middle of the heatmaps bins. As *S*_0_ is negative *E*_2_ is negative also, as this is a linear we can simply take the absolute value of *E*_2_. For the Threshold of Binocularity, *E*_2_ represents the rate of decrease, with eccentricity, of the maximum spatial wavelength at which binocular processing can occur.

### A wavelength-eccentricity surface on binocular thresholds

The ToB as calculated from our ICA resembles a sigmoid function. In order to extend our analysis to consider both eccentricity and wavelength we need to define a sigmoid function over two-dimensions. We define a probability *p*(*β|μ, λ*) of a sample component taken from eccentricity *μ* and wavelength that the component is binocular. The probability that the component is monocular is then 1 *- P* (*β|μ, λ*). The standard sigmoid can be extended to a two-dimensional form as:

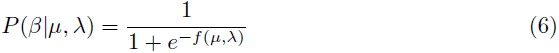

Where *f* is a simple linear function

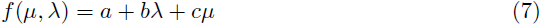

The free parameters *a*,*b*,& *c* were determined by fitting *p*(*β|μ, λ*) to our data using least squares minimisation tools from Matlab’s curve fitting toolbox.

The ToB is the iso-contour of the 2D sigmoid function where the value *p*(*β|μ, λ*) = 0.5.

The 2D sigmoid forms 1D sigmoids in both the wavelength and eccentricity dimensions (to see this hold either *λ* or *μ* constant). The threshold function can be directly calculated from the linear equation 5 (to see this set equation 6 equal to 0.5). *E*_2_ as defined in equation 4 is a linear equation, in this form it cannot be calculated from a sigmoid. However, an alternative definition of *E*_2_ defines *E*_2_ as the ‘eccentricity at which [the stimulus size] is twice the foveal value’ [29]. In our case the sigmoid is at its maximum at the fovea rather than its minimum, therefore we invert the concept and find the eccentricity at which the threshold of binocularity is half that at the fovea. This concept is shown in diagrammatic form in figure 7.

**Figure 7.**
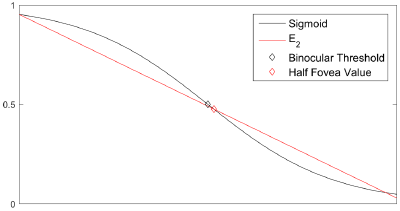
*E*_2_ on the sigmoid.

Here *E*_2_ is defined as the value on the sigmoid that is half the foveal value. *E*_2_ is now related to the gradient of a linear function between the value of the sigmoid at the fovea (0) and the eccentricity at which the sigmoid is half the value of the fovea. Clearly if the value of the sigmoid at the fovea is 1 then *E*_2_ will be equal to the eccentric at the binocular threshold where the value of the sigmoid is 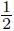. In the example above the sigmoid value never reaches unit (and common occurrence in our data), thus the Half Fovea value is not exactly equal to the binocular threshold (0.5).

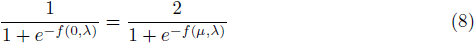

We can calculate *E*_2_ by solving equation 8 for *λ*. Here the left hand side returns the value of the sigmoid at 0 eccentricity (*μ* = 0), and the right hand side is twice the value of the unmodified sigmoid. Equation 8 can be solved for *μ* as:

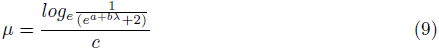

Clearly when the binocular/monocular ratio at the fovea is one, *E*_2_ is equal to the threshold of binocularity and can be found trivially using equation 9. As the value at the fovea tends to 1, the value of equation 9 will tend to equation 7.

## Supporting Information

Binocular photographic image data and (Matlab) source code associated with this publication is available from GitHub at https://github.com/DavidWilliamHunter/Bivis.

## 1 Acknowledgements

This work was supported the Biotechnology and Biological Sciences Research Council [grant number BB/K018973/1]

## Author contributions

Conceived and designed the experiments: DWH PBH. Performed the experiments: DWH. Analyzed the data: DWH. Contributed reagents/materials/analysis tools: PBH. Wrote the paper: DWH PBH.

## Additional Information

The author(s) declare no competing financial interests.

